# Bisphenol TMC disturbs mitochondrial activity and biogenesis, reducing lifespan and healthspan in the nematode *Caenorhabditis elegans*

**DOI:** 10.1101/2024.05.20.595050

**Authors:** Laxmi Rathor, Ho Jeong Lee, Taylor McElroy, Steven Beck, Julia Bailey, Stephanie Wohlgemuth, Sung-Hwan Kim, Jeong-doo Heo, Rui Xiao, Sung Min Han, Moonjung Hyun

**Author notes:** Co-corresponding authors: Moonjung Hyun, Sung Min Han. Co-first authors (Laxmi Rathor, Ho Jeong Lee, Taylor McElroy).

## Abstract

Rising concerns about Bisphenol A (BPA) toxicity have prompted the search for safer alternatives. However, concerns persist regarding the safety of replacements like bisphenol TMC (BPTMC), a rapidly emerging BPA substitute. Utilizing the *in vivo* model organism *Caenorhabditis elegans* (*C. elegans*), whose shared genes mirror human biology, we aim to unveil the potential toxicity of BPTMC on a live animal. *C. elegans* exposed to 1 mM BPTMC exhibited developmental delays, reduced reproduction, and diminished longevity. Furthermore, an investigation into mortality at various animal stages, oxidative stress, and thermal stress revealed additional compromised toxicity. Notably, exposure to BPTMC resulted in mitochondrial abnormalities, including reduced oxygen consumption, lowered mitochondrial membrane potential, and decreased ATP levels. Additionally, BPTMC increased ROS levels but decreased mitochondrial population. Transcriptome analysis revealed that BPTMC induces alterations in the expression of genes associated with mitochondrial biogenesis. Our findings raise crucial concerns about BPTMC as a safe BPA alternative. The observed widespread toxicity across key life stages suggests a need for further investigation into the potential toxicity of BPTMC on human health and environmental consequences.

## 1. Introduction

The widespread use of plastics in our daily lives is undeniable, yet their potential health consequences remain a source of growing concern. Among the synthetic polymers raising suspicion is BPA, a ubiquitous monomer employed in the production of food containers, water bottles, dental supplies, canned food, baby bottles, and even receipt paper ^1, 2^. It is imperative to recognize that BPA, owing to its structural resemblance to the female hormone estrogen, falls under the category of endocrine disruptors, eliciting its influence by binding itself to estrogen receptors in the human body, thus operating analogous to a hormone ^3–5^. The ever-increasing acknowledgment of the potential health risks posed by BPA has necessitated the exploration and development of alternative compounds like BPS, BPF, and, more recently bisphenol BPTMC. However, this pursuit of safer alternatives has not dispelled worries, as the safety profiles of these substitutes remain shrouded in ambiguity ^5^.

BPTMC, similar to BPA, has been detected in household goods like food packaging and baby bottles, further highlighting its presence in our daily lives ^6^. Recent studies have revealed the presence of BPTMC in materials employed for food packaging. The investigation of the migration of bisphenols from baby bottles from 20 brands reveals that BPTMC has the fourth largest detection frequency (55%) among 14 tested bisphenols including BPA ^7^. Additionally, another study to determine the predominant bisphenols in Norwegian sewage sludge and sediment and Czech surface water shows that BPTMC frequently found compounds in sediment water ^8^.

Due to its structural similarity to other harmful bisphenols and its growing presence in everyday products, further research on BPTMC has emerged to assess its potential toxicity. Several recent studies have highlighted the potential endocrine-disrupting and toxic effects of BPTMC ^9, 10^. Recent research in *Oryzias melastigma*, a fish species, demonstrates detrimental developmental, behavioral, and endocrine effects following exposure to the chemical compound BPTMC ^9^. Notably, BPTMC exposure has been shown to delay hatching, increase aggressive behavior, and alter estrogen receptor binding. Furthermore, results from both in silico and in vitro experiments using *Saccharomyces cerevisiae* suggest that BPTMC strongly binds to the estrogen receptor ^9^. Additionally, exposure to BPTMC has been shown to stimulate testosterone secretion in the culture of the rat fetal testis, suggesting potential endocrine-disrupting activity ^10^. *In vitro*, assays also report that BPTMC can affect lipid droplet numbers in cultured MA-10 and KGN cells ^11^. Moreover, it has been noted that BPTMC exhibits stronger anti-progestogenic activities compared to BPA ^8^. Despite these initial findings, our understanding of BPTMC toxicity *in vivo* remains incomplete. Given the widespread presence of BPTMC in everyday products and the potential for human health consequences, these findings raise concerns about the safety of BPTMC as a replacement for BPA and emphasize the need for further research to elucidate its complete toxicological profile.

*C. elegans* shares many similar genes with humans, making it a valuable model organism for studying diseases and understanding how molecules work. Given our previous discovery that other bisphenol compounds like BPS, BPF, and TMBPF harm *C. elegans* by affecting lifespan, reproduction, and survival, we further investigated the potential impact of BPTMC using this powerful *in vivo* model ^12^. Exposure to 1 mM BPTMC resulted in significant developmental delays, reduced reproduction, and diminished longevity in *C. elegans*. Additionally, increased sensitivity to oxidative and thermal stress was observed, along with mitochondrial abnormalities, including reduced oxygen consumption, lowered mitochondrial membrane potential, and decreased ATP levels. Moreover, BPTMC exposure led to an increase in reactive oxygen species (ROS) levels while decreasing mitochondrial population. Finally, transcriptome analysis unveiled alterations in the expression of genes associated with mitochondrial biogenesis, suggesting the potential impact of BPTMC on mitochondrial homeostasis at the molecular level. This study underscores the urgency of comprehensively assessing the safety and impact of BPTMC, given its widespread use in various consumer goods and the potential risks it may pose to human health and the environment.

## 2. Materials and Methods

### 2.1. Chemicals

Dimethyl sulfoxide (DMSO, 99.7% pure), Bisphenol TMC (Catalog #, 52746-49-3) and 2′,7′-Dichlorodihydrofluorescein diacetate (Catalog #, D6883) were purchased from Millipore Sig ma (MO, USA) and Santa Cruz Biotechnology (Catalog #, sc-503712A; TX, USA). MitoTrack er Red CMXRos (Catalog #, M46752) and MitoTracker Green FM (Catalog #, M7514) were purchased from ThermoFisher Scientific (MA, USA).

### 2.2. C. elegans

*C. elegans* strains, including the wild-type (WT): N2, CF1553: muIs84 [(pAD76) sod-3P::GFP + rol-6(su1006)], TJ356: zcIs356 [daf-16p::daf-16a/b::GFP + rol-6(su1006)], SJ4100: zcIs13 [hsp-6p::GFP + lin-15(+)] and PE255: feIs5 [sur-5p::luciferase::GFP + rol-6 (su1006)], were maintained at 20 °C on NGM plates seeded with *Escherichia coli* (*E. coli*) OP50 ^13^. Chemical plates, containing BPTMC and DMSO at a final concentration of 1mM, were prepared and dissolved in DMSO. Control plates with DMSO alone were also created. The plates were left overnight at ∼22 °C for chemical equilibration into the agar. All strains were obtained from the Caenorhabditis Genetics Center (CGC).

### 2.3. Lifespan assay

About 25-30 synchronized animals at the L4 larval stage were picked individually and transferred to fresh NGM plates containing DMSO alone as control or 1 mM BPTMC. The animals were transferred to new DMSO or BPTMC NGM plates every three days. Each day, animal mortality was tested based on their response to a platinum wire touch. All experiments were independently conducted at least five times at 20 °C.

### 2.4. Development assay

Five 1-day-old-stage WT N2 were picked from NGM and transferred individually to NGM plates treated with DMSO or 1 mM BPTMC. After 3 hours, the animals were removed, and synchronized embryos were continuously cultured. After 3 days (about 65 hours later), the developmental stages of the animals were scored. At least 100∼200 animals for the developmental stage were tested for each group. All experiments were conducted at 20 °C and performed in triplicate.

### 2.5. Reproduction assay

Three WT N2 larvae at the L4 stage were picked up from normal NGM and individually transferred to a fresh OP50 seeded NGM plate treated with DMSO or 1 mM BPTMC (2∼3 each plate). The number of laid embryos was scored under a microscope. The experiment was repeated daily until no eggs were laid. At least three independent experiments were performed for each condition.

### 2.6. Lethality assay

Worms were prepared and synchronized by egg preparation. For egg lethality assay, eggs were transferred to DMSO as a control or with 1 mM BPTMC in 1 ml M9 buffer. For the L1 lethality assay, synchronized L1 larval animals were transferred to DMSO as a control or with 1 mM BPTMC in 1 mL M9 buffer. For adult lethality assay, synchronized 1-day-old adults were transferred to DMSO as a control or with 1 mM BPTMC in 1 mL M9 buffer. For the viability assay after 24 hours of treatment, the animals were transferred to a fresh NGM plate, and the response to the platinum wire touch was scored. At least three independent experiments were performed for each condition.

### 2.7. Stress tolerance assay

For oxidative stress analysis, synchronized L4 larval animals were transferred to each NGM plate containing DMSO or 1 mM BPTMC. After 24 hours, 1-day-old adults were harvested from each condition and transferred individually to 12 well plates (30-50 worm/well) that contained 0, 0.1, and 0.2 mM H_2_O_2_ in 1 mL M9 buffer. The animals were then incubated at 20 °C for 24 hours. For the viability assay, the animals were transferred to a fresh NGM plate, and the response to the platinum wire touch was scored. At least three independent experiments were performed for each condition. For UV irradiation assay, 1-day-old animals treated with DMSO and 1mM BPTMC were exposed to a dose of 1000 J/m^2^ in a UV chamber. The percentage survival was then checked after 48 hours and 72 hours. For thermal stress assay, worms were prepared and synchronized by egg preparation. Synchronized L4 larval stages were transferred individually to each NGM OP50 seeded plate containing DMSO or 1 mM BPTMC, and incubated at 20 °C. After 24 hours, for the thermal stress assay, 20-30 synchronized 1-day-old animals were incubated for 5 hours at 35 °C and then incubated again overnight at 20 °C. They were recorded as dead when there was no response to contact with a platinum wire.

### 2.8. MitoTracker staining

MitoTracker Red CMXRos and Green FM were utilized *in vivo* to assess mitochondrial membrane potential and mass in 3-day-old WT N2 adults ^14^. Following 1mM BPTMC or DMSO treatment from the L1 stage, about 50 L4 larvae were transferred to new OP50-seeded NGM plates containing 100 µM MitoTracker. After 2 days, worms were moved to normal OP50 seeded NGM plates, cultured in the dark for 1 hour, and washed with M9 buffer. Images were taken with a spinning disk confocal microscope, and fluorescence intensity was measured using ImageJ. Each condition underwent at least three independent experiments.

### 2.9. Oxygen consumption rate in *C. elegans*

Oxygen consumption rate (OCR) was measured using a Seahorse XFe96 Analyzer (Agilent) following a previously described protocol ^15, 16^. Age-synchronized L1 stage worms were cultured on NGM agar plates containing 1 mM BPTMC for 72 hours, then transferred to fresh NGM plates with the same BPTMC concentration. After 2 days, worms were harvested and washed with M9 buffer. 15 worms per well in M9 buffer (200 microliter) were transferred to a Seahorse XF96 V3 PS Cell Culture Microplate (Agilent) with 10 technical replicates per treatment. OCR was measured as follows: basal OCR in five cycles, maximal uncoupled OCR after FCCP injection (10 µM), and residual non-mitochondrial OCR (ROX) after sodium azide injection (40 mM). The number of worms in each well was confirmed at the end of the assay with a Cytation-1 Imaging Multi-Mode Reader (BioTek Instruments). Data were analyzed using Agilent/Seahorse software Wave Desktop and Controller (v2.6.1) and Microsoft Excel, presenting ROX-corrected OCR per worm (pmolO2/min/worm; mean ± standard error of the mean of 10 technical replicates).

### 2.10. Detection of ROS in *C. elegans*

ROS levels were measured with H_2_DCFDA dye dissolved in DMSO to make 10mM stock and kept in dark condition. For the staining, synchronized wild-type N2 L4 larval pre-treated with BPTMC and DMSO were transferred into a new OP50 seeded chemicals plate for a day at 20 degrees. 1-day-old WT N2 worms were moved from 20 degrees to 35 degrees for 1 hour. After 1 hour, worms were washed with M9 buffer three times to remove bacteria and resuspended in M9 to a final volume of 500 µl. 2.5µl of H_2_DCFDA solution was added to a final concentration of 50µM. Worms were incubated at room temperature (∼22 °C) for 1 hour in a dark condition in a shaking incubator. After 1 hour, worms were again washed three times in M9 buffer and transferred to a 2% agarose slide and covered with a cover slip.

Whole body fluorescence intensity was measured with Image J. For the spectrophotometry, Wild-type N2 worms were maintained on NGM plates containing DMSO or BPTMC from L1 larval to the 1-day-old adult stage at 20 °C and washed with M9 buffer to remove bacteria. ∼150 1-day-old worms from the control group and BPTMC were transferred into 96 well plates and added 1.5 µl of H_2_DCFDA from 10 mM stock solution into each well to make a 50 µM final concentration in a 300 µl volume. The observation was recorded for 140 min at intervals of 20 min in a fluorescence spectrophotometer at the excitation wavelength of 485 nm and the emission wavelength of 528 nm at room temperature.

### 2.11. Quantification of stress response pathways in *C. elegans*

Age-synchronized populations of CF1553, SJ4100, and TJ356 strains were obtained for *sod-3p*::GFP, P*hsp-6*::GFP, and DAF-16::GFP expressions, respectively. Initially, worms at the L1 stage were transferred to an OP50-seeded NGM plate containing either 1mM BPTMC or DMSO alone. Upon reaching the L4 stage, the worms were transferred to a fresh chemical plate. On the second day, 25-30 worms were utilized for imaging. The fluorescence intensity of the worms was observed using a fluorescence microscope system (Nikon ECLIPSE Ti) and Crest X-Light-V3 spinning disk confocal microscope equipped with Nikon TiE and Prime 95B 25 mm camera (Crest Optics, Nikon, and Photometrics, respectively). Subsequently, the fluorescence intensity of each worm was measured using ImageJ and Nikon Elements.

### 2.12. Neuronal morphology in *C. elegans*

A well-established transgenic line was used to analyze neuronal morphology that expresses soluble GFP in GABAergic neurons (*juIs76*). L1 stage transgenic animals were grown on OP50-seeded NGM plate containing DMSO or 1mM BPTMC chemicals until the L4 stage was attained, after which they were transferred to fresh NGM plates treated with BPTMC. After 2 days, the GABAergic motor neurons in BPTMC-treated worms were imaged to assess the number of beadings in an individual neurite.

### 2.13. *C. elegans* mobility assay

Body size and mobility assays were performed using an automated worm tracker instrument, providing an unbiased analysis (WormLab, MBF Bioscience, USA) ^17^. WT N2 worms, synchronized at the L1 stage, were cultured on either DMSO or 1mM BPTMC NGM plates supplemented with OP50 at a controlled temperature of 20°C. Upon reaching the L4 developmental stage, the worms were transferred to fresh chemical NGM plates pre-seeded with OP50 and allowed to age until reaching 3-day-old adult. Subsequently, on the second day of adulthood, the worms were placed beneath a worm tracker instrument, and a video was recorded for 30 seconds utilizing a Basler acA2500 camera, coupled with a 10x close-focus zoom lens to minimize potential light-induced stress. Behavioral parameters, encompassing body speed, length, and wavelength, were assessed using, were assessed using the MDF MSCOP-010 animal lab IL laminator base and WormLab software.

### 2.14. Transcriptome analysis

#### 2.14.1. RNA isolation

Total RNA was isolated utilizing Trizol reagent (Invitrogen). The evaluation of RNA quality was conducted utilizing the Agilent TapeStation 4000 system (Agilent Technologies, Amstelveen, The Netherlands), while quantification of RNA was achieved employing the ND-2000 Spectrophotometer (Thermo Inc., DE, USA).

#### 2.14.2. Library preparation and sequencing

The QuantSeq 3’ mRNA-seq Library Prep Kit (Lexogen, Inc., Austria) was used to build libraries for control and test RNAs, following the manufacturer’s instructions. In brief, each total RNA was prepared, and an oligo-dT primer with an Illumina-compatible sequence at the 5’ end was hybridized to the RNA before reverse transcription was performed. Following the degradation of the RNA template, a random primer with an Illumina-compatible linker sequence at the 5’ end was used to initiate second-strand synthesis. The double-stranded library was purified using magnetic beads to remove all reaction components. The library was expanded to include the complete adapter sequences needed for cluster generation. The finished library is purified using PCR components. NextSeq 550 (Illumina, Inc., USA) was utilized for high throughput sequencing, specifically single-end 75 sequencings.

#### 2.14.3. Data analysis

The QuantSeq 3’ mRNA-seq readings were aligned using Bowtie2 (Langmead and Salzberg, 2012). Bowtie2 indices were created using genome assembly or representative transcript sequences for genome and transcriptome alignment. The alignment file was used to assemble transcripts, estimate their abundances, and identify gene expression differences. Counts from unique and multiple alignments were used to find differentially expressed genes using Bedtools (Quinlan AR, 2010). The RC (Read Count) data were normalized using the TMM+CPM approach using EdgeR within R (R development Core Team, 2020) and Bioconductor (Gentleman et al., 2004). DAVID (http://david.abcc.ncifcrf.gov/) and Medline database (http://www.ncbi.nlm.nih.gov/) searches were used to determine gene categorization. ExDEGA (Ebiogen Inc., Korea) was used for data mining and graph visualization.

## 3. Results

### 3.1. Effect of BPTMC exposure on *C. elegans* development and lethality

Early exposure to BPA could interrupt normal growth and development ^12, 18^. The life cycle of *C. elegans* consists of an embryonic stage, four larval stages (L1-L4), and adulthood^19^. To understand the impact of BPTMC exposure on the development and growth of animals, we cultured worms at each growth stage (egg, L1, and adult) in liquid M9 buffer containing 1 mM BPTMC for 24 hours and evaluated survival rates (Figure 1A). When the harvested eggs grown on normal condition were transferred and incubated in M9 buffer containing 1 mM BPTMC, 100% of them failed to hatch in the BPTMC-treated group, while the control group showed minimal lethality (3.4%) (Figure 1B). This difference strongly suggests that BPTMC possesses the capacity to penetrate through eggshells, thereby exerting a profound effect on embryonic development. Additionally, the L1 stage worms, exposure to BPTMC, exhibited 91% lethality, a dramatic increase from the 0% observed in the control group (Figure 1C). This enhanced lethality persisted into the post-developmental stage, affecting the lethality of adult animals (Control: 0.5% vs. BPTMC: 100%) (Figure 1D). Intrigued by these results, we further investigate the developmental state of worms cultured on solid NGM containing 1 mM BPTMC (BPTMC-NGM) for 65 hours post-egg laying. While exposing L1 stage worms to 1 mM BPTMC in M9 buffer resulted in complete lethality, a significant portion of worms cultured on 1 mM BPTMC-NGM successfully reached the L4 or adult stage (Figure 1E). This suggests that worms were likely exposed to less BPTMC in the BPTMC-NGM condition compared to the BPTMC-M9 liquid buffer condition. However, when compared to the DMSO-NGM control condition, where the majority of worms reached the adult stage (approximately 88%) after 65 hours post-egg laying, only 27% of worms cultured on the BPTMC-NGM developed into adulthood (Figure 1E). We used the BPTMC-NGM condition for the subsequent experiments. Next, considering the previously reported adverse effect of BPS on body growth in *C. elegans*, we also examined the body growth of worms exposed to BPTMC by measuring the body length and width using an automated WormLab in free-moving status^20^. The body length and width of the C. elegans were significantly reduced when they were cultured on a 1 mM BPTMC-NGM compared with the DMSO-NGM control (Figures 1F and G). The total body dimensions assessed by imaging mounted worms on slides decreased with age, suggesting reduced body growth (Figure 1H). Finally, we investigated the impact of BPTMC on reproduction by counting the number of embryos laid by *C. elegans* exposed to BPTMC. Our observations revealed a decrease in the average number of embryos in the cultured group on BPTMC-NGM (254.5 per worm) compared to the control group (325.8 per worm) (Figure 1I). Consequently, BPTMC demonstrated an unequivocal capacity to elevate lethality rates across all developmental stages, from the germline to adulthood.

**Figure 1.**
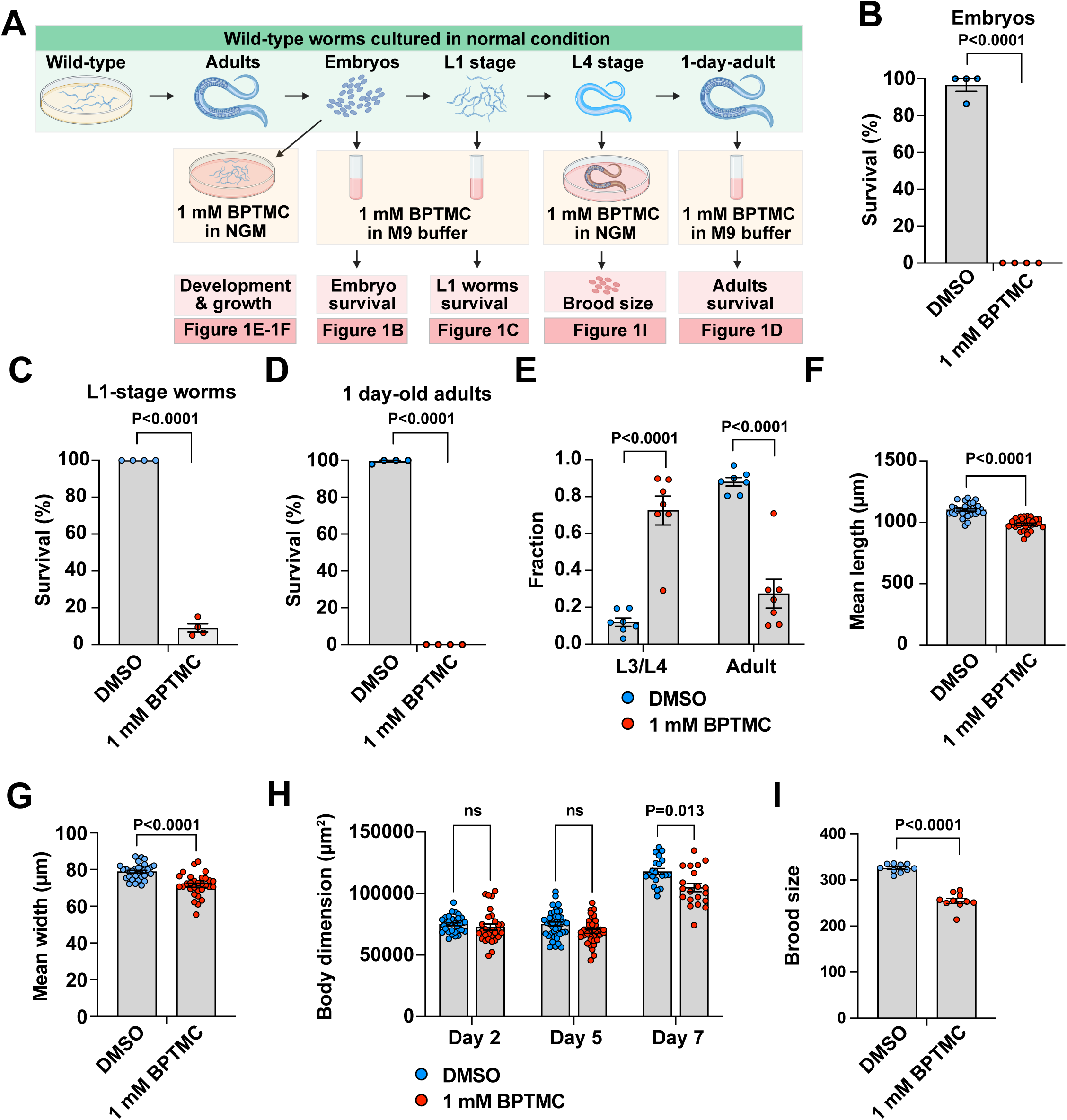
Effect of BPTMC on the lethality, development, and reproduction. (A) A diagram illustrating lethality, development, and reproduction experimental procedures is provided. We measured the survival rate of worms at specified stages after incubating them in a liquid M9 buffer containing 1 mM BPTMC or DMSO for 24 hours. The brood size was evaluated by transferring L4 stage worms grown on normal NGM conditions to NGM containing either DMSO only or 1 mM BPTMC. (B-D) The survival after exposure to BPTMC for 24 hours in each development stage of embryo (B), L1 (C), and adult (D). (E) The fraction of adulthood 65 hours after eggs were released, and the number of each developmental stage were scored. (F-H) Body growth parameters at 2-day-old adults including mean length (F), mean width (G), and body dimension (H). (I) The average number of eggs laid. Individual dots indicate a single set (B, C, D, E, and I) or individual worm (F, G, and H). The values are presented as mean ± SEM. p-value indicates the difference with the DMSO-treated control group. Two-tailed student’s t-test (B, C, and D); Two-way ANOVA (E and H); two-tailed Mann–Whitney test (F, G, and I).

### 3.2. Effect of exposure to BPTMC on the lifespan and aging of *C. elegans*

To examine the impact of BPTMC exposure on the lifespan of worms, we synchronized L4 stage worms cultured under basal conditions and transferred and maintained them throughout their lifespan on control DMSO-NGM and 1 mM BPTMC-NGM. The control group showed a lifespan, achieving a 50% survival rate of 12.71 days. In contrast, exposure to BPTMC resulted in a remarkable reduction in lifespan, with only 50% of the worms surviving to 4.33 days (Figure 2A). This marked discrepancy represents a significant decrease of approximately 66% (P.<0.0001) in lifespan following exposure to BPTMC. To determine whether the reduced lifespan of BPTMC-treated worms was due to accelerated aging or to sickness, symptoms of normal aging, such as lipofuscin accumulation, were investigated in worms exposed to BPTMC ^21^. Notably, the level of lipofuscin, an aging hallmark that typically increases progressively over time, was decreased in 1-day-old adult worms treated with BPTMC compared to the control group ^21^ (Figures 2B and 2C). This lipofuscin level was significantly increased at the 5-day-old stage, suggesting faster aging in BPTMC-exposed worms until the mid-age point (Figures 2B and 2C). In line with the findings from a previous report indicating a decrease in lipofuscin levels approximately 6–12 hours before death, 7-day-old BPTMC-treated worms, showing high mortality rates in the lifespan assay (Figure 2A), exhibited a reduction in the level of lipofuscin (Figures 2B and 2C) ^22^. These observations suggest the adverse impact of BPTMC on the aging process.

**Figure 2.**
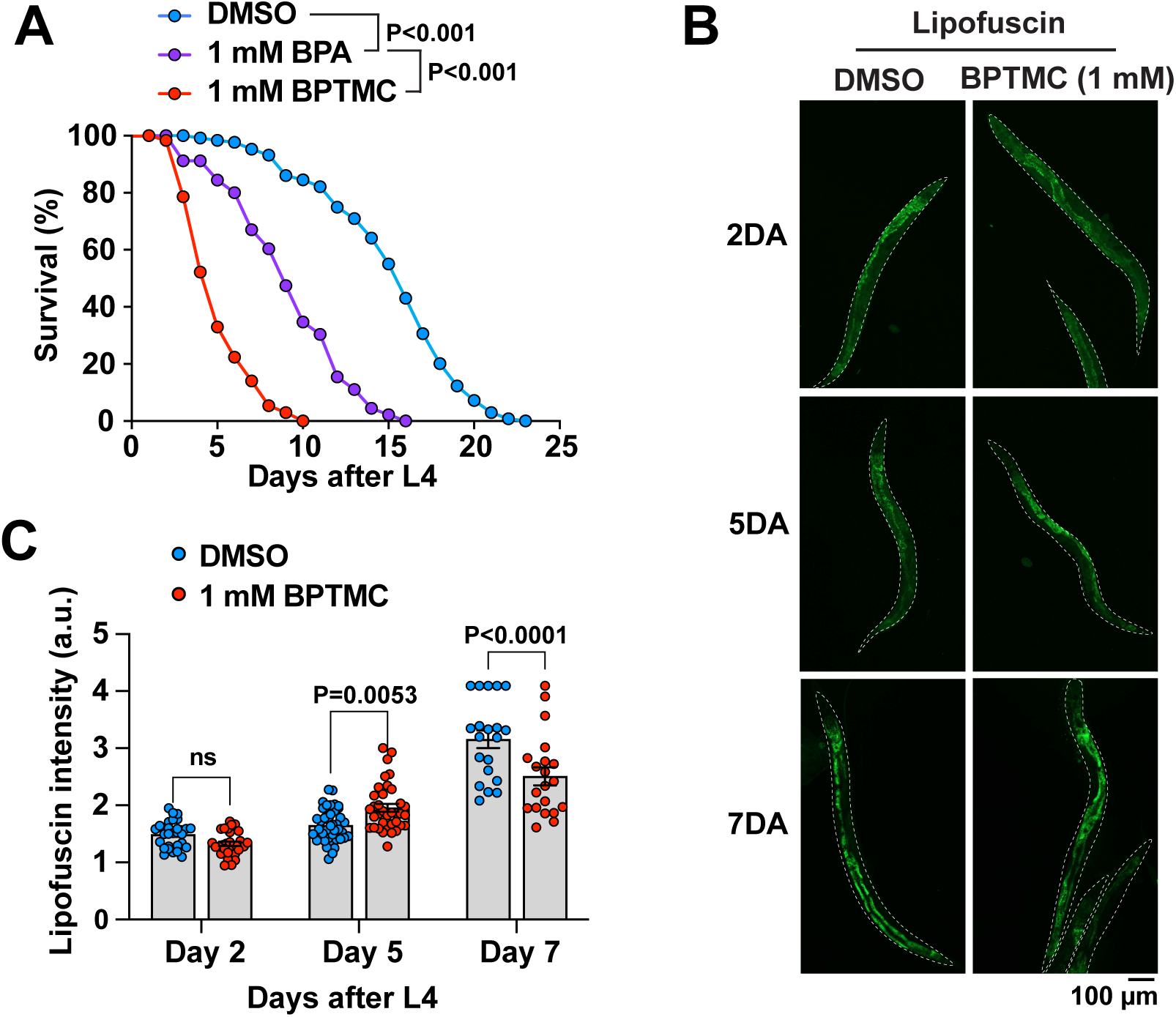
Effect of BPTMC on lifespan and aging. (A) The survival curve represents the average of at least 5 sets of results for each different condition. The significance of the lifespan curves was assessed using a Log-rank (Mantel-Cox) test. (B) The representative images of lipofuscin fluorescence indicated days after the L4 stage. (C) Quantification of lipofuscin intensity. Each dot indicates a single worm. The values are presented as mean ± SEM. Two-way ANOVA.

### 3.3. Effect of exposure to BPTMC on the neuronal aging of *C. elegans*

Bisphenols have detrimental effects on the nervous system ^23–25^. In addition, organismal aging is heterogeneous, with each tissue, such as the nervous system, exhibiting a unique aging rate at the cellular level ^26–30^. The occurrence of swelling or beading in neurites has been recognized as a hallmark of aging ^27, 31, 32^. Additionally, neuronal beading is associated with neurodegenerative diseases and has been observed across a diverse range of species, from *C. elegans* to mammals ^27, 31^. We explore the effects of BPTMC on the beading formation in GABAergic motor neurons that form commissural neurites connecting the dorsal and ventral nerve cords ^33^ (Figure 3A). In the 2-day-old adult group, the BPTMC treatment significantly increased the number of beadings within the neurite compared to the control group (p=0.03, n=60 axons for each group) (Figures 3B and 3C). Additionally, the dorsal nerve cord also exhibited increased beading formation (Figure 3B). Our results suggest that BPTMC could accelerate aging-related changes in the neurons.

**Figure 3.**
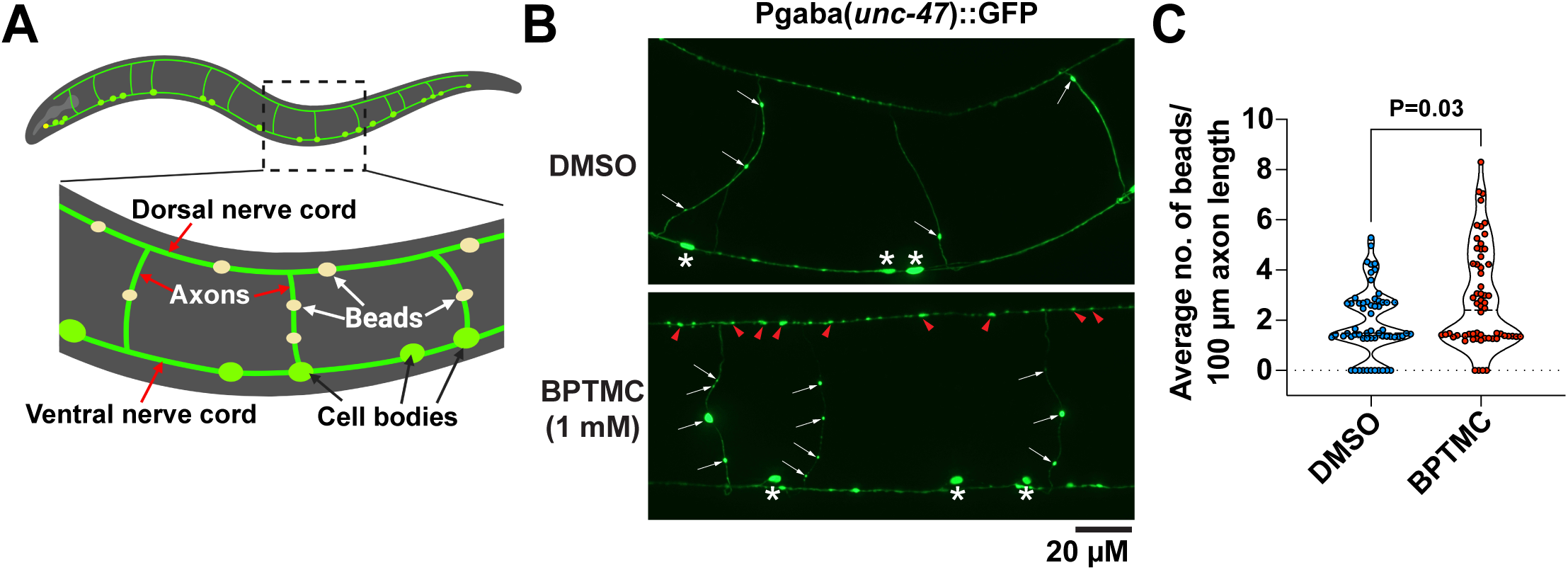
Effect of BPTMC on neuronal aging. (A) Diagram and representative images of the beads formed within *C. elegans* GABAergic motor neurons which extend commissural neurites from the ventral nerve cord to the dorsal nerve cord. Each neurite is extended from a single neuron. (B and C) Representative image (B) and quantification (C) of the beads within a single axon of 2-day-old worms. (B) Arrows indicate beads on commissural axons. Arrowheads indicate bead structures on the dorsal nerve cord. Asterisks indicate the cell bodies of GABAergic motor neurons on the ventral nerve cord. (C) Each dot indicates the bead number within a single axon. The values are presented as mean ± SEM. Two-tailed Mann–Whitney test.

### 3.4 Effect of exposure to BPTMC on the healthspan of *C. elegans*

Lifespan is associated with healthspan, but not in all cases ^34–36^. We thus investigated the impact of BPTMC exposure on various healthspan parameters in *C. elegans*, including body motility and tolerance to stressors ^35–42^. First, we analyzed the effect of BPTMC on resistance to thermal and oxidative stresses, which are healthspan parameters closely linked to longevity across species ^43–47^. Adult worms from both the control and BPTMC groups were treated with varying doses of hydrogen peroxide (H_2_O_2_) for 24 hours. As a result, BPTMC exposure significantly decreased survival rate in a dose-dependent manner compared to the control group (Figure 4A). Furthermore, worms were subjected to UV irradiation, a major source of DNA damage and accelerating aging ^48–50^. The BPTMC-exposed group exhibited a roughly 41% decrease in survival compared to the control group (Figure 4B). Next, worms from both groups were exposed to a 35 °C environment for 5 hours to test their heat stress tolerance. We found a reduced survival rate to thermal stress within the BPTMC-exposed group compared to the control (Figure 4C). Next, we measured the body motion parameters on solid media, including crawling speed and the wave-like body bending posture formed during crawling, using segmented images recorded with an automated worm behavior tracking system ^29, 34, 37, 40^ (Figure 4D). While exposure to 1 mM BPTMC did not alter the maximal velocity of body crawling, it significantly decreased the average crawling speed (Figures 4E and 4F). In line with this result, the wavelength (λ) of the body wave normalized by body length was decreased by 1 mM BPTMC exposure (Figure 4G). Together, these results indicate that BPTMC exposure affects the healthspan of *C. elegans* at various degrees depending on the parameters.

**Figure 4.**
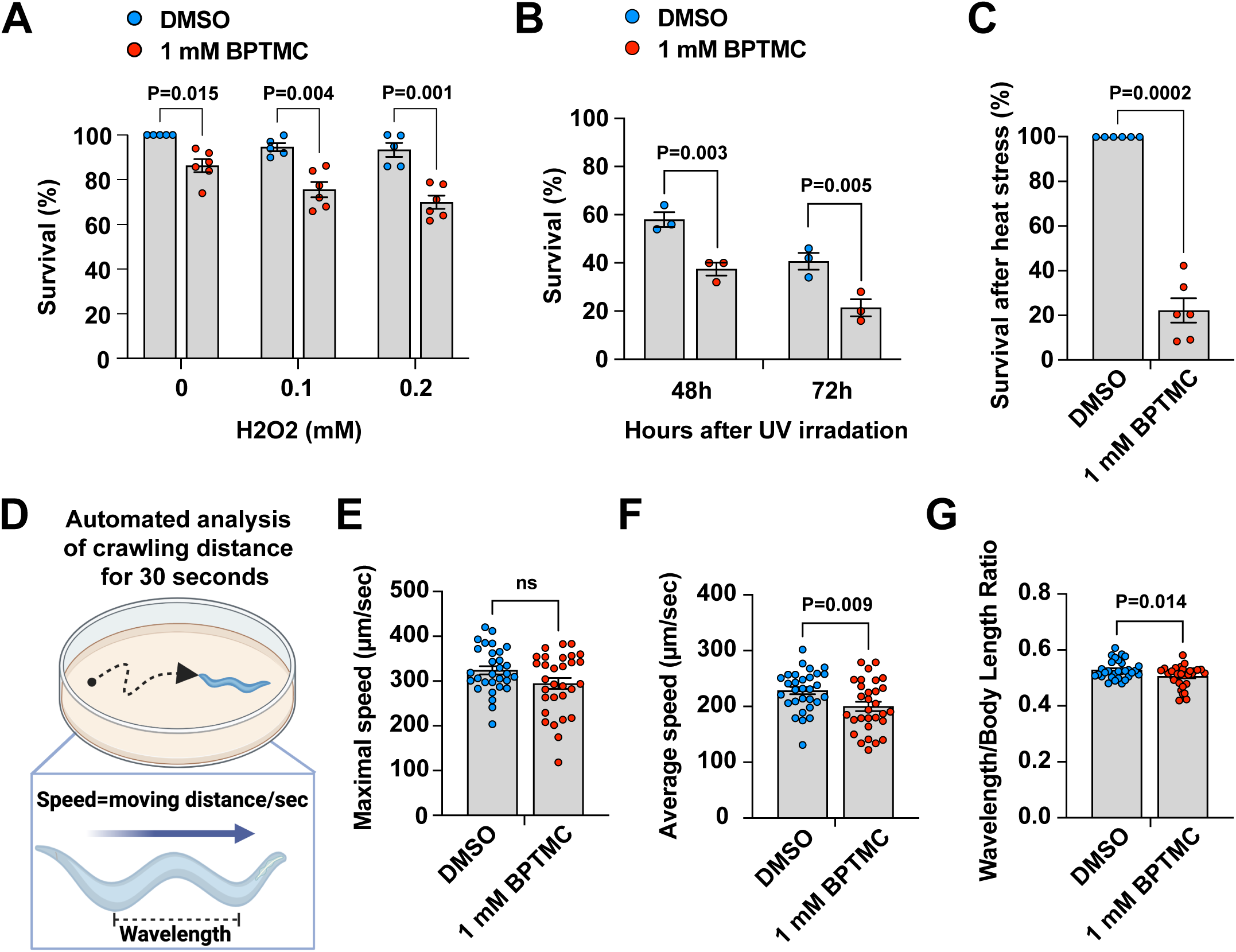
Effect of BPTMC on healthspan parameters. (A) Survival rate of animals cultured in the indicated concentration of H_2_O_2_. (B and C) The survival rate after UV irradiation (B) and incubation at 35°C for 5 hours (C). For A, B, and C, approximately 30 to 50 1-day-old adults are tested for each set. (D) Illustration of track length and wavelength measurements. (E) Maximal crawling, (F) average crawling speed, and (G) body-bending wavelength. The track length is recorded for 30 seconds. Each dot indicates an individual tested worm’s value. The values are presented as mean ± SEM. Two-way ANOVA (A and B); two-tailed Mann–Whitney test (C, E, F, and G).

### 3.5. Effect of BPTMC on mitochondrial activity

Multiple lines of evidence indicate the implication of oxidative stress and mitochondrial dysfunction in the adverse health outcomes attributed to BPA ^51–54^. Additionally, mitochondrial defects are a key determinant of aging ^55, 56^. To test the impact of BPTMC on mitochondrial homeostasis, we subjected L1 stage *C. elegans* to BPTMC exposure until they reached 2-day-old adulthood. Mitochondrial membrane potential, a key indicator of mitochondrial function, was assessed using MitoTracker CMXRos, a red-fluorescent dye sensitive to the membrane potential ^57, 58^. Intriguingly, BPTMC significantly diminished mitochondrial membrane potential, evident from the reduction in red fluorescence intensity (Figures 5A and 5B). This observation was further verified by Seahorse XF extracellular flux analysis. We found that BPTMC exposure decreased the overall oxygen consumption rate (Figure 5C). Both basal and maximal oxygen consumption rates in BPTMC-exposed groups were markedly reduced compared to the control group (Figures 5D and 5E). These results indicated that exposure to BPTMC impaired the function of mitochondria in a live animal.

**Figure 5.**
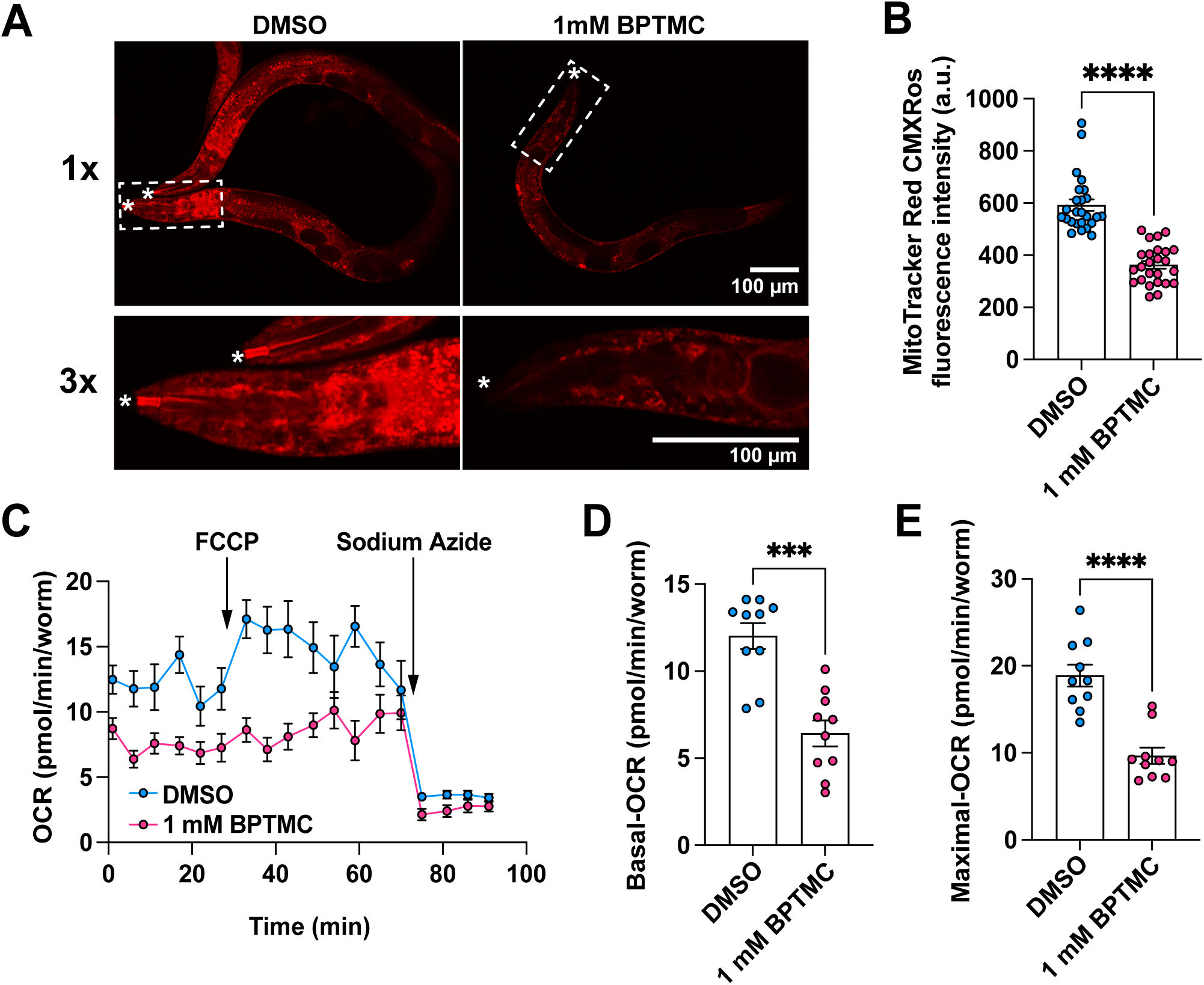
Effect of BPTMC on mitochondrial function. (A and B) Representative images (A) and quantification (B) of mitochondrial membrane potential after treatment with BPTMC. MitoTracker CMXRos is used to determine mitochondrial membrane potential. In (A), dashed box areas are magnified below. Asterisks indicate the tip of the head. In (B), each dot indicates a single worm. (C-E) The Seahorse Bioscience XF24 extracellular flux analyzer was also employed to measure basal and maximal OCR (pmol/min/worm), indicating OXPHOS activity. (C) Schematic of Seahorse mitochondrial oxygen consumption analysis. Following the establishment of a baseline, sequential addition of FCCP and sodium azide was performed. (D) The basal OCR was determined by calculating the difference between the mean OCR values during the baseline period and during sodium azide treatment. (E) The maximal OCR was obtained by calculating the difference between the mean OCR values during the FCCP treatment and during sodium azide treatment. Each dot indicates a single set consisting of 15 worms. The values are presented as mean ± SEM. Two-tailed Mann–Whitney test.

### 3.6. Effect of BPTMC on the ROS level and stress response pathway

Intrigued by the alterations in mitochondrial functions in worms treated with BPTMC, we further explore the influence of BPTMC on mitochondrial integrity by evaluating reactive oxygen species (ROS) levels within the worms. The worms were treated with BPTMC from the L1 stage to day 1 of adulthood and then stained with H2DCF-DA ^59, 60^. ROS level evaluated by imaging DCF fluorescence was increased in all BPTMC-treated groups compared to the control group (Figures 6A and 6B). Consistently, DCF-fluorescence accessed by the spectrophotometer of a group of worms supports the increased ROS levels by BPTMC exposure (Figure 6C). We then tested whether BPTMC enhances antioxidant mechanisms, including superoxide 3 (SOD-3, an ortholog of human SOD2) ^61^. To explore this, we utilized an integrated transgenic line expressing GFP under the control of the *sod-3* promoter (P*sod-3*::GFP) ^62, 63^. P*sod-3*::GFP expression levels were monitored at least 18 hours after exposure to BPTMC. As a result, *sod-3* expression was significantly elevated throughout the bodies of worms exposed to BPTMC compared to the DMSO-treated controls (Figures 6D and 6E). Forkhead transcription factor DAF-16/FoxO factor plays a key role in modulating *C. elegans* lifespan and the expression of *sod-3* ^64–66^. We conducted additional experiments to assess the influence of BPTMC on DAF-16/FoxO activity by examining the localization of DAF-16::GFP. Normally, DAF-16 proteins are primarily located in the cytoplasm, but they translocate to the nucleus when exposed to stress ^64–66^. Worms exposed to BPTMC exhibited no notable increase in the translocation of DAF-16::GFP to the nucleus compared to control (Figures 6F and 6G). The mitochondrial unfolded protein response (mitoUPR) is another pathway associated with mitochondrial stress and its activation can be evaluated by monitoring the well-characterized *zcls13[hsp-6p::gfp]* mitoUPR reporter line ^67^. Interestingly, BPTMC exposure enhanced mitoUPR activation, as evidenced by increased expression of P*hsp-6*::GFP (Figures 6I and 6J). Together, these results suggest that exposure to BPTMC increases ROS levels, leading to the induction of mitoUPR in worms. However, it does not activate DAF-16/FOXO as a protective response against ROS.

**Figure 6.**
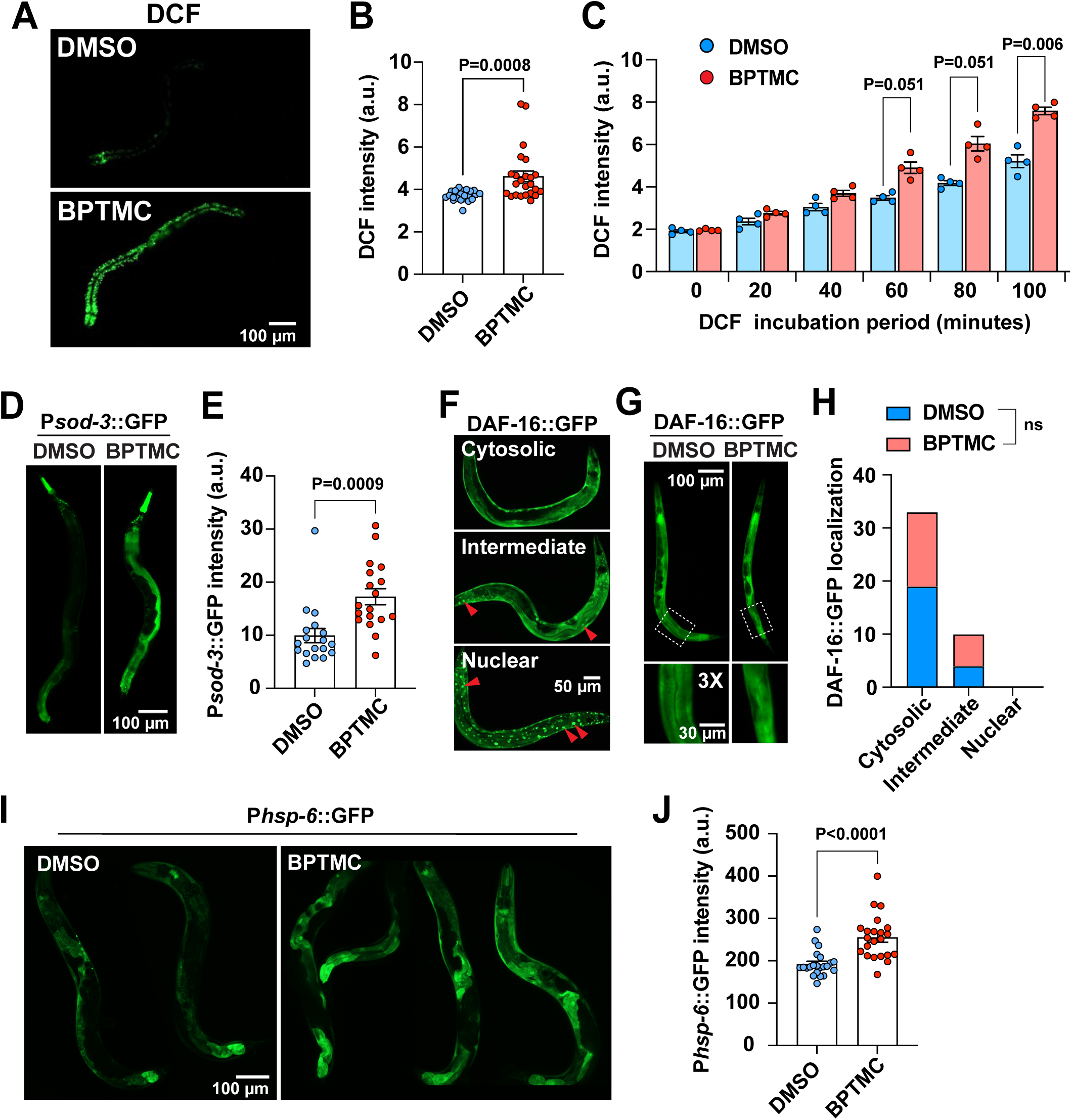
Effect of BPTMC on ROS homeostasis. (A and B) Representative images (A) and quantification (B) of ROS levels measured by DCF fluorescence intensity. Each dot represents the value of an individual animal. (C) Time-dependent DCF intensity increase was assessed using spectrophotometry over the indicated time window after DCF staining. Each dot represents a single set of the assay with 150 animals at the 1-day-old adult stage. E) Representative images of P*sod-3*::GFP expression after BPTMC exposure. (E) Quantification of P*sod-3*::GFP expression. (F) Representative images of DAF-16::GFP expression in cytoplasm, intermediate, and nucleus. (G and H) Representative images (G) and quantification (H) of DAF-16::GFP expression after BPTMC exposure. DAF-16::GFP was mainly observed in the cytoplasm in both DMSO control and BPTMC-treated groups. For each group, approximately 20–24 animals were tested. (I and J) Representative images (I) and quantification (J) of P*hsp-6*::GFP expression after BPTMC exposure. Each dot represents a single worm. For all graphs, values are presented as means ± SEM. The p-value indicates the difference with DMSO-treated control; Two-tailed Mann–Whitney test (B, E, and J); Two-way ANOVA (C); Chi-square test (H).

### 3.7. Effect of BPTMC on the expression of mitochondrial homeostasis-associated genes

It has been documented that exposure to BPA results in a significant reduction in mitochondrial biogenesis and related protein levels ^68, 69^. Our data also indicated the decreased mitochondrial mass and DNA copy number, suggesting a potential impact of BPTMC on mitochondrial biogenesis. We conducted a transcriptome analysis to evaluate the expression of critical genes associated with mitochondrial biogenesis that may be altered by BPTMC exposure. Two replicates of DMSO (control group) and BPTMC-treated (1 mM) worms were subjected to sequencing (Figure 7A). Our results indicated that 32 up-regulated differently expressed genes (DEGs) (Table S1) and 90 down-regulated DEGs (Table S2) following BPTMC exposure (fold change of 1.5 with a p-value < 0.05) (Figure 7B). To identify the key biological processes influenced by BPTMC exposure, DEG annotation, and enrichment analysis were carried out to dissect the distribution of DEGs across GO categories, which are sub-categorized as molecular function (MF), biological process (BP), and cellular component (CC) subsets (Carbon et al., 2017). A total of 16 significantly enriched GO terms were identified. Among these, 4 terms related to MF, 7 to BP, and 5 to CC (Table S3, Figure 7C). Subsequently, the detailed sub-GO terms of each subset were refined to evaluate the overall impact of BPTMC (Figures 7D-7F). Prominent among the key GO terms, most significantly enriched in each category, were those related to mitochondrial biogenesis, including BP-GO:0032543 (mitochondrial translation), MF-GO:003735 (a structural constituent of ribosome), and CC-GO:005763 (mitochondrial small ribosomal subunit). Notable genes enriched within these GO terms encompass *mrps-16/mrps16*, *mrps-34/mrps34*, *mrpl-44/mrpl44*, and *dap-3/dap3*. These genes are predicted to encode highly conserved mitochondrial ribosomal proteins crucial for translating mitochondrial DNA-encoded genes, required for mitochondrial biogenesis ^70^ (Figures 7B and 7G). Also, the expression of *F53G2.12*, *pde-12/pde12,* and *C38B6.5/gatb* was decreased. These genes are predicted be associated with mitochondrial biogenesis by regulating the iron-sulfur cluster assembly in ETC ^71^, polyadenylation of mitochondrial mRNA ^72^, and glutaminyl-tRNA synthase ^73^, respectively. In line with the increased ROS level, an increase in expression was noted for *icl-1*, a gene pivotal in protecting worms against heightened mitochondrial ROS levels by facilitating an effective activation of the mitochondrial unfolded protein response (mitoUPR) (Figures 7B and 7G) ^74^. To further assess the impact of BPTMC on mitochondrial biogenesis, we examined the copy number of mitochondrial DNA. Our findings revealed a decrease in mitochondrial DNA copy number in the BPTMC-exposed group compared to the control group (Figure 7H). Consistent with these results, the overall mitochondrial mass across the body, as evaluated by staining with MitoTracker Green FM, was significantly reduced in the BPTMC-treated groups (Figure 7I) ^75^. Collectively, these transcriptome data revealed significant alterations in gene expression profiles and enrichment of key biological processes, particularly those associated with mitochondrial translation and ribosomal function, suggesting potential mechanisms underlying BPTMC-induced effects on mitochondrial biogenesis.

**Figure 7.**
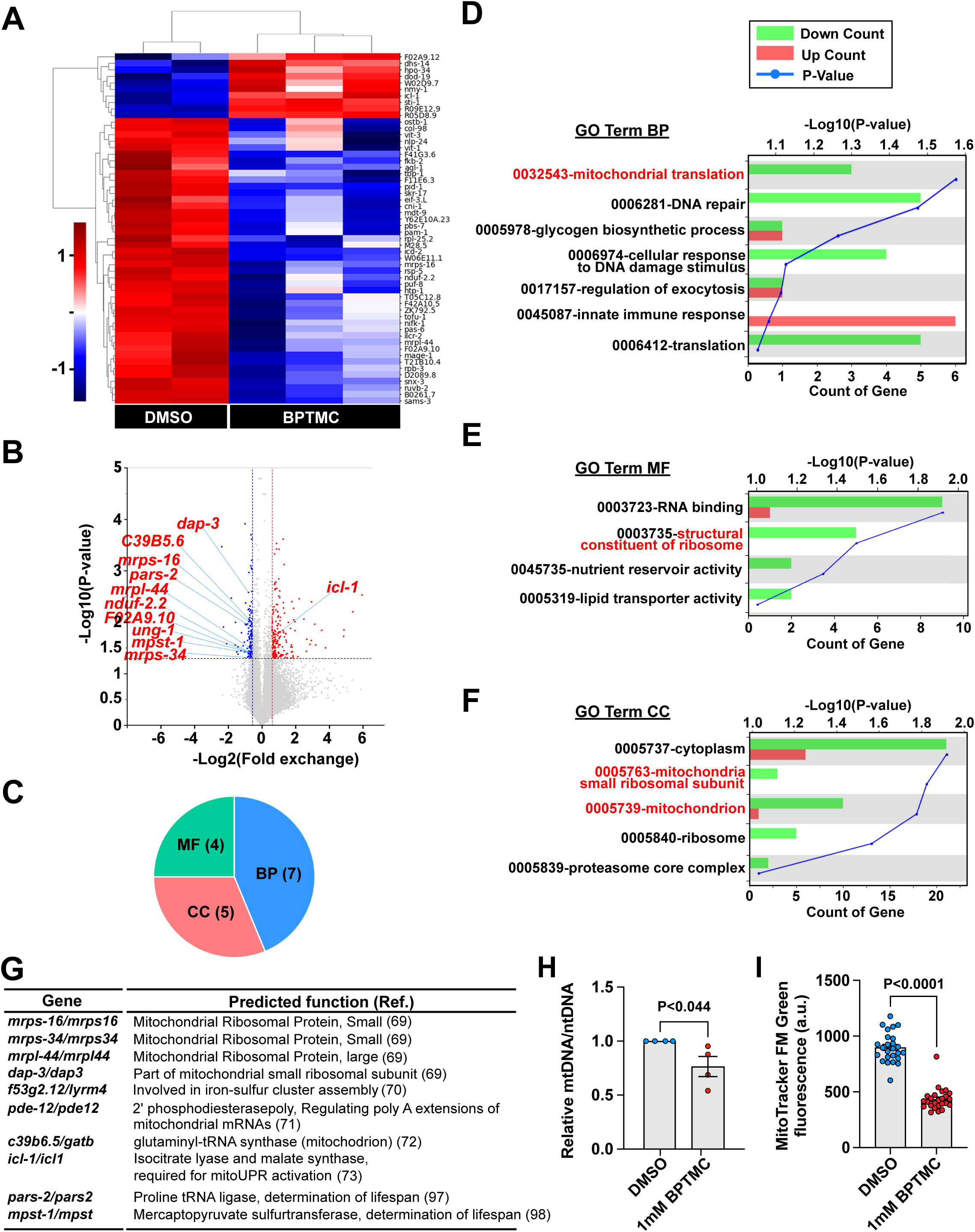
The transcriptome changes induced by BPTMC exposure. (A) Heatmap displaying two replicates of DMSO-treated (control group) and three replicates of BPTMC-treated worms that were sequenced. (B) Volcano plots comparing the control groups with the BMTMC-treated group. (C) Pie chart showing the number of significantly enriched GO terms. (D-F) GO term classifications into three subsets: molecular function (MF) (D), biological process (BP) (E), and cellular component (CC) (F). (G) List of key DEG genes associated with mitochondria. (H) qPCR analysis of mitochondrial genome copy number assessed by ND1 encoded in mtDNA and normalized by COX-1 encoded in the nuclear genome. The values are presented as means ± SEM. (I) Relative MitoTracker FM Green fluorescence intensity of the bisphenol-treated group compared to that in the DMSO control group. Each dot represents an individual animal’s value. Two-tailed Student’s t-test (H); Two-tailed Mann– Whitney test (I).

## 4. Discussion

The widespread use of BPA in various products and its ability to invade the human body through diverse pathways emphasize the urgent need to explore safer alternatives. Studies report a significant prevalence of BPA in human urine samples, maternal breast milk, and even in healthy infants with no apparent history of exposure ^76–78^. Given the documented adverse effects of BPA on human health, including endocrine disruption, cancer, hepatotoxicity, neurodevelopment, and cognitive abilities, the need for alternatives like BPTMC is evident and necessitates investigation ^79–84^.

Environmental stress stands out as a significant determinant influencing nearly every aspect of life events in organisms such as *C. elegans* ^85^. Despite the inherent stress resilience of *C. elegans*, exposure to environmental chemicals, such as bisphenol, can compromise this resilience, resulting in adverse consequences such as defective development and reduced healthspan and lifespan. Our findings further support the adverse impact of BPTMC on worm development, reproduction, and aging parameters, including lifespan and healthspan. Similar to BPTMC, BPA exposure in *C. elegans* has been linked to the acceleration of the aging process and a shortened lifespan ^86^. Our previous studies in *C. elegans* also highlight the impact of multiple BPA analogs on diminishing the lifespan ^12^. However, bisphenol impacts on lifespan vary since recent investigations involving marine rotifers and *C. elegans* show that BPS has no discernible effect on lifespan, whereas BPA decreased it ^87^. Our current study showed a dramatic 66% reduction in lifespan in the BPTMC-exposed group compared to the control group. This reduction is among the largest observed in worms exposed to BPA or other bisphenols in our previous report ^12^. Importantly, the diminished lifespan was accompanied by a high degree of lethality across all stages, from embryos to the adult stage. Additionally, the observed reduction in reproductive ability suggests consistent defects in germline development, making BPTMC a critical toxicity factor in both developmental and post-developmental stages throughout the lifespan of an animal.

Numerous lines of evidence support the crucial role of mitochondria in the aging process, as their dysfunction impacts both lifespan and healthspan ^55, 56^. The nature of these changes depends on the type and degree of mitochondrial dysfunction and ROS level, leading to either enhanced or decreased parameters of lifespan and healthspan, indicating a complex regulatory role of mitochondria in aging ^88, 89^. Our finding showing the toxic impact of BPTMC on mitochondria suggests its potential role in regulating health and longevity.

Previous research has linked BPA exposure to mitochondrial damage and oxidative stress in both human and worm model systems ^53, 90^ ^91–94^. For instance, BPA exposure decreases the activities of mitochondrial ETC complexes in rat liver ^53^. Our prior research in *C. elegans* uncovered that BPA and various BPA analogs induce a reduction in mitochondrial membrane potential, oxygen consumption, and mitochondrial mass ^12^. Intriguingly, each analog exhibits a distinct impact on mitochondrial homeostasis. For example, TMBPF decreases mitochondrial mass in *C. elegans* but does not significantly alter oxygen consumption rate, reactive oxygen species levels, and membrane potential, indicating intricate effects of different bisphenols on mitochondrial health ^12^. In cultured mammalian cells, exposure to BPA and its analogs, such as TMBPF, BPS, and others, results in increased mitochondrial ROS levels ^12^. This highlights the broad disruption of mitochondria by bisphenols. Our current study extends this exploration to BPTMC, revealing a decrease in mitochondrial membrane potential and oxygen consumption rate, suggesting that mitochondria are likely a major target broadly disrupted by bisphenols, including BPTMC. Importantly, exposure to BPTMC resulted in a decrease in mitochondrial mass and mitochondrial DNA copy number, suggesting a potential association with reduced energy production. The reduced mitochondrial DNA copy numbers have also been reported in human CD8□T cells after long-term exposure to low-dose BPA ^69^. Intriguingly, worms exposed to BPTMC exhibited significantly increased ROS levels despite a reduction in mitochondrial mass. This suggests not only diminished mitochondrial mass but also the presence of mitochondrial defects, which contribute to increased ROS formation.

Transcriptome analysis also further illuminated the molecular consequences of BPTMC exposure. BPTMC exposure decreased the expression of 7 genes, encoding nuclear genes related to mitochondrial ribosomal proteins and mitochondrial mRNA and tRNA processing, which play a pivotal role in protein synthesis within the mitochondrion ^71–73^. This observed decrement in these genes suggests potential disruptions in mitochondrial protein synthesis, compromising the assembly of mitochondrial ribosomes and overall mitochondrial function. Notably, BPA inhibits mitochondrial biogenesis by impairing mitochondrial protein import in the rat brain hippocampus, suggesting that bisphenols could broadly affect the mitochondrial protein transport process ^68^. Furthermore, BPTMC exposure resulted in decreased expression of *pars-2* and *mpst-1*. Notably, depletion in these genes results in reduced lifespan in *C. elegans* ^95, 96^.

Our results indicate that despite the reduced mitochondrial mass and activity, BPTMC increases ROS levels. Defects in mitochondrial electron transport chains are often associated with increased ROS levels, along with a reduced membrane potential, suggesting that BPTMC could damage mitochondrial quality ^56, 97^. Although its protective role remains unknown, our previous study in *C. elegans* demonstrated that exposure to BPS significantly upregulated the expression of *sod-3*, a well-known antioxidant enzyme, and its upstream regulator DAF-16/FoxO transcription factor ^12, 64–66^. Our current study found that exposure to BPTMC increased *sod-3* expression but did not alter DAF-16/FoxO activity, suggesting that BPTMC led to both defective mitochondria producing higher ROS levels and reduced protective activity. This could account for the dramatic reduction in lifespan and survival of worms exposed to BPTMC. Alternatively, these results suggest the activation of alternative protective mechanisms against mitochondrial defects caused by BPTMC. In line with this view, we found that BPTMC enhanced mitoUPR activity. Furthermore, our transcriptome analysis showed an increase in *icl-1* expression. This gene encodes a sole protein that catalyzes the glyoxylate shunt in *C. elegans* ^74^. A recent study indicates that ICL-1 also plays a critical role in triggering mitoUPR in response to elevated levels of mitochondrial ROS ^74^. This suggests that worms may utilize the mitoUPR pathway rather than the DAF-16/FoxO pathway to protect themselves against the increase in ROS associated with BPTMC exposure. The functional roles of these mechanisms require further study. Additionally, the impact of BPTMC on other mitochondrial stress response pathways, such as the mitochondrial unfolded protein response, remains a subject for further investigation.

Finally, our study utilized 1 mM concentration of BPTMC, mirroring those in previous research on BPA and other bisphenols but potentially exceeding typical environmental levels ^51, 98–100^. Notably, recent studies, employing a new testing method, disclosed markedly higher total BPA levels (51.99 ng/ml) in urine samples from pregnant women compared to the previously suggested mean value (1.24 ng/ml) reported by the National Health and Nutrition Examination Survey ^76, 77^. The significant difference observed prompts a reconsideration of total bisphenol levels in humans, including the BPTMC level. Additionally, *C. elegans* exhibits significant resistance to pharmacologically active molecules, requiring much higher concentrations for effects compared to mammalian cell cultures. This resistance is due to the extensive xenobiotic defenses of worms, which include their tough cuticle and xenobiotic efflux pumps. These defenses act as physical barriers and rapidly expel xenobiotics, significantly reducing the amount of absorbed chemicals reaching the body ^85, 101–104^. Moreover, our findings indicate that the effects observed under the 1 mM BPTMC-NGM condition were considerably weaker compared to those observed under the 1 mM BPTMC-M9 liquid buffer condition. This suggests that the treatment of worms by culturing on solid media containing BPTMC is likely to reduce the actual exposure concentration. Hence, the actual concentration of BPTMC absorbed by worms could be substantially lower than the treated concentration.

In conclusion, our comprehensive exploration highlights the complex impact of BPTMC on mitochondrial function and gene expression in *C. elegans*. The harmful effects on mitochondrial dynamics, biogenesis, and the expression of mitochondria-related genes underscore the intricate interplay between BPTMC exposures and mitochondrial homeostasis. These findings contribute valuable insights to the broader understanding of the implications of bisphenol exposure on cellular and molecular processes, emphasizing the need for further research to elucidate specific mechanisms and potential protective pathways.

## Acknowledgment

Some strains were provided by the CGC, funded by the NIH Office of Research Infrastructure Programs (P40 OD010440). This research was conducted while SMH was a Hevolution/AFAR New Investigator Awardee in Aging Biology and Geroscience Research (AGR00030264), and the Korea Institute of Toxicology (KIT; KK-2406-05 to J.D.H), the National Research Foundation of Korea (NRF, 2021R1F1A1045599, 2022R1F1A1069374 to M.J.H).

## Author Contributions

LR performed mitochondria and stress tolerance experiments. TM performed mobility and neuronal aging experiments. HJL performed lifespan, lethality, development, and healthspan experiments. SW Performed oxygen consumption experiments. SB performed the analysis of lipofuscin-related data. JB prepared reagents and assisted in experimental procedures. RX, SHK, and JDH provided counseling and advice on experimental design and analysis. HJL, LR, SMH, and MH contributed to writing the manuscript. SMH and MJH contributed equally to supervising the research. All authors discussed the results and contributed to the final manuscript.

